# Human generation times across the past 250,000 years

**DOI:** 10.1101/2021.09.07.459333

**Authors:** Richard J. Wang, Samer I. Al-Saffar, Jeffrey Rogers, Matthew W. Hahn

## Abstract

The generation times of our recent ancestors can tell us about both the biology and social organization of prehistoric humans, placing human evolution on an absolute timescale. We present a method for predicting historic male and female generation times based on changes in the mutation spectrum. Our analyses of whole-genome data reveal an average generation time of 26.9 years across the past 250,000 years, with fathers consistently older (30.7 years) than mothers (23.2 years). Shifts in sex-averaged generation times have been driven primarily by changes to the age of paternity rather than maternity, though we report a disproportionate increase in female generation times over the past several thousand years. We also find a large difference in generation times among populations, with samples from current African populations showing longer ancestral generation times than non-Africans for over a hundred thousand years, reaching back to a time when all humans occupied Africa.

## Main text

Knowledge of the human generation time (or “generation interval”) in the recent past is important for many fields. While genetic data has provided deep insights into human history, population genetic methods typically scale history in terms of generations (e.g. *1, 2*). This makes knowing the generation time especially important for determining the absolute timing of historic events, including migrations to new continents (*3*) or gene flow with extinct hominids (*4*). In order to transform these population genetic estimates into absolute time, it is commonly assumed that current generation times have persisted across hundreds of thousands of years, or that studies of extant hunter-gatherer (forager) societies provide representative generation times across the span of human history (*5, 6*). However, neither assumption is likely to be correct: the average age at which males and females have children depends on many environmental, demographic, and cultural factors that can change rapidly (*7*), while contemporary hunter-gatherer societies differ substantially from each other and from past societies (*8*). It is also clear that generation times have evolved among the great apes (*9*), and may therefore have evolved along the branch leading to modern humans.

Previous genetic approaches to estimating historical generation times (the average age at which individuals conceive children) have taken advantage of the compounding effects of either recombination (*10*) or mutation (*11*) on modern human DNA sequence divergence from ancient samples. While these estimates have provided significant insight, they are averaged both across the sexes and across the past 40-45 thousand years. Greater resolution is possible by examining the mutations that originated at specific times in the past, together with a model that accurately predicts the generation times of individuals producing those mutations. Here, we develop a model that uses the spectrum of *de novo* mutations as a predictor of parental age. By coupling this model with variants whose ages have been estimated from genome-wide genealogical information, we are able to separately estimate the male and female generation times at many different points across the past 250,000 years.

As humans age, the number and type of *de novo* mutations they transmit to their offspring changes (*12, 13*). We use information on mutations from a large pedigree study with parents whose ages at conception are known (*14*) to model the relationship between parental age and the counts of the six different types of single-nucleotide mutations (Fig. S1). The mutation counts are modeled as a multinomial draw from a distribution with a probability vector that is itself drawn from a Dirichlet distribution (Supplemental Methods); we regress mutation counts on both paternal and maternal age in this Dirichlet-multinomial model (Fig. 1A; Fig. S2). After filtering mutations using the same criteria applied to segregating variants (see below), our model was trained on 27,902 phased mutations from 1,247 trios. To obtain mutation spectra from many different periods in the past, we used the estimated time of origin for current polymorphisms from the Genealogical Estimation of Variant Age (GEVA) approach (Fig. 1B; ref. (*15*)). This method estimates when in the past each of ~43 million variants from the 1000 Genomes Project arose by mutation. We excluded variants from mutation classes that may have been multiply mapped on genealogies (e.g., CpG→TpG mutations), as well as TCC→TTC transitions, which have been inferred to be the result of a recent mutation pulse (*16, 17*). After filtering, we retained 25.3 million variants. More than 80% of the sampled variants arose in the last 10,000 generations, but very few are from the last 100 generations (Fig. S4). Because the sampled variants are so unevenly distributed through time, we divided the data from the past 10,000 generations into bins with equal numbers of variants. Maternal and paternal ages were estimated by fitting the variant spectrum in each of these bins to our Dirichlet-multinomial model by minimizing compositional (Aitchison) distance between the observed spectrum and the model (Supplemental Methods). This set of parental ages form the estimate of the generation interval.

**Figure 1.**
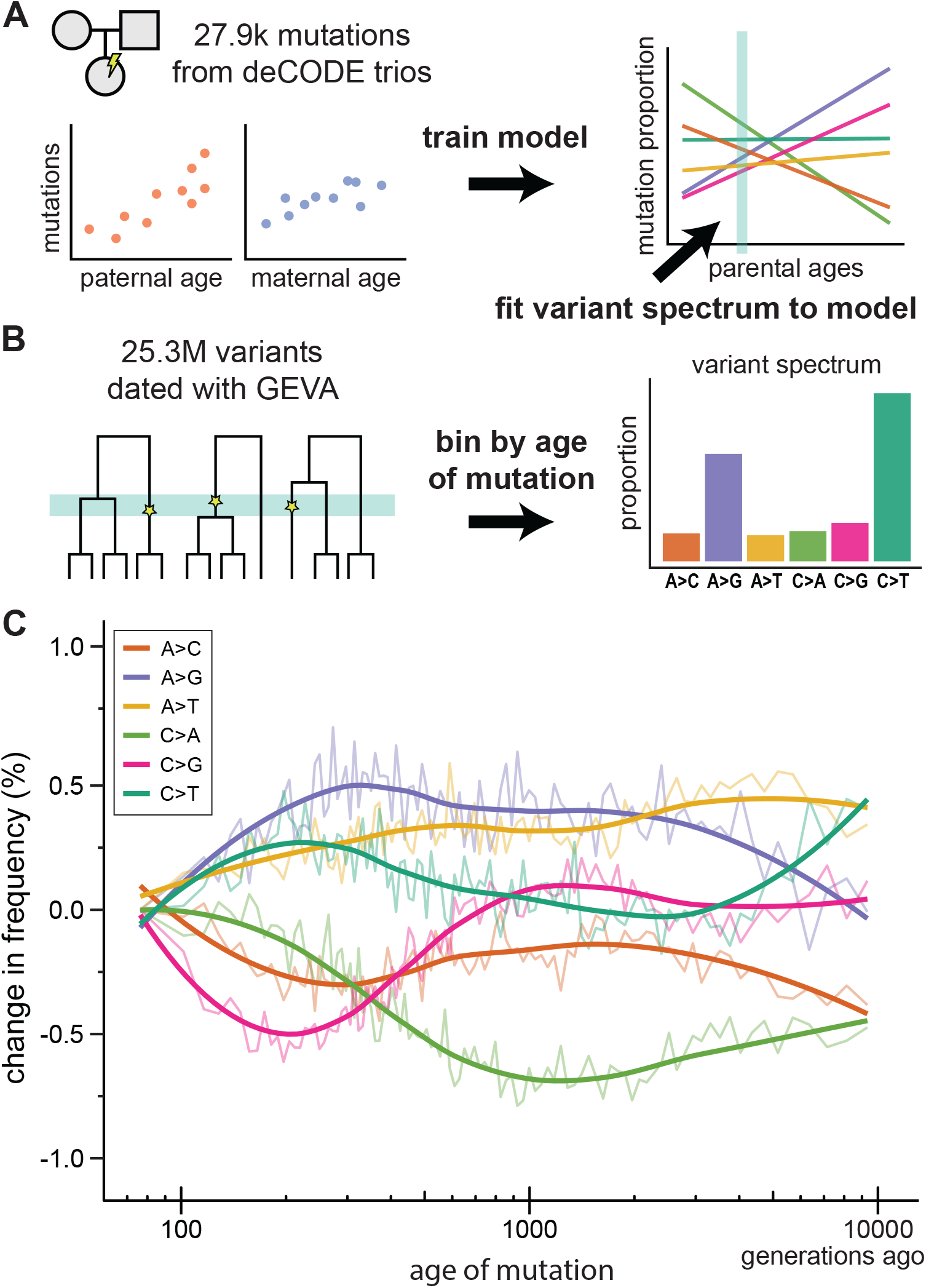
The mutation spectrum changes with human generation time. (A) Data on *de novo* mutations from 1,247 Icelandic trios (*14*) were used to train a model that predicts the effect of both maternal and paternal age on the mutation spectrum. (B) Data from 25.3 million segregating variants whose date of origin was estimated using GEVA (*15*) were used to assess the mutation spectrum at different periods in the past. The mutation spectrum from each time period (bin) was used as input to the model from panel A in order to estimate the generation interval for males and females. (C) Differences in the frequency of each of the six different mutation types through time, as compared to the most recent time period (smoothed lines from local regression). Fig. S3 presents the absolute frequencies of the same mutation data over time.

These data allow us to estimate generation times for males and females across the past 250,000 years (Fig. 2A). Within this timeframe, we find the average human generation interval to be 26.9 ± 3.4 years (standard error) with an average for males of 30.7 ± 4.8 years and an average for females of 23.2 ± 3.0 years. The results show that human generation times have undergone a rapid increase in the recent past after declining for over a thousand generations. The average human generation interval was at a recent minimum of 24.9 ± 3.5 years at ~250 generations ago (6.4 kya), roughly concurrent with the historic rise of early civilizations. Before this, it had declined from a peak of 29.8 ± 4.1 years at ~1,400 generations ago (38 kya), just before the beginning of the Last Glacial Maximum. The trends found for human generation time were not affected by the stringency of mutation filters on the training set, nor by the effects of either biased gene conversion or linked selection on the variant spectrum (Fig. S5, S6).

**Figure 2.**
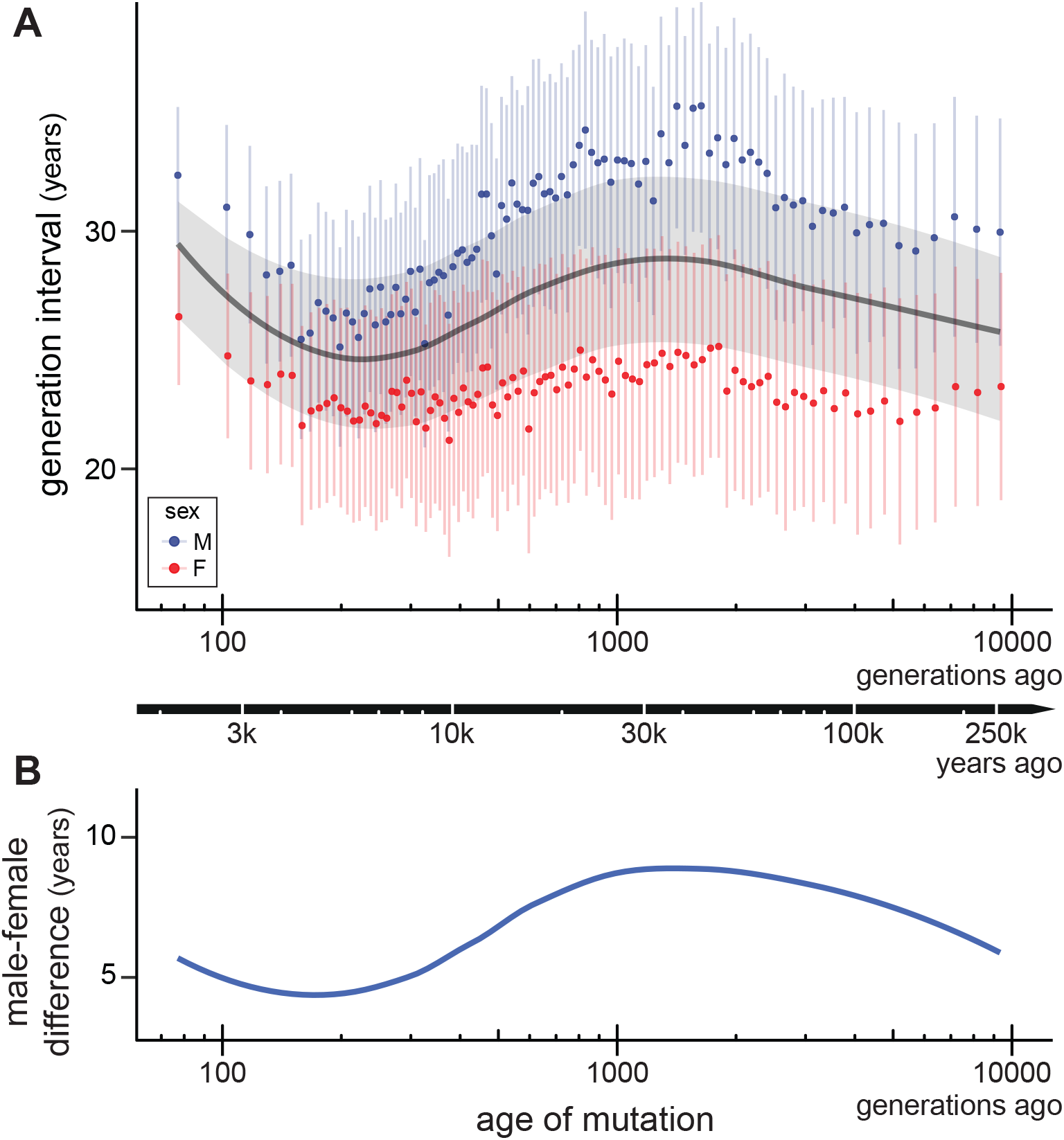
Estimating the male and female generation interval across 250,000 years. (A) Male (blue points), female (red points), and sex-averaged (grey line) generation intervals over the past 10,000 generations. The data were divided into 100 time periods with equal numbers of variants; generation intervals in each were independently estimated using the Dirichlet-multinomial model. Sex-averaged generation intervals are shown here as a line smoothed by local regression. Confidence intervals (±1 S.E.) displayed for all estimates were obtained by resampling both the *de novo* mutation data for bootstrapped models and the variants in each time period for bootstrapped spectra. The absolute timeline (black arrow) was calculated by integrating sex-averaged generation-time estimates across generations elapsed since the present (Supplementary Methods). (B) The smoothed difference (loess) between estimates of the male and female generation interval over time.

Our model estimates a longer generation interval for males than females across all analyzed time periods (Fig. 2B). These results are consistent with studies of contemporary cultures, more than 99% of which show a longer male generation interval (*5*). Overall, there is a high correlation between the average generation interval and the male-female difference (Pearson’s *r* = 0.88; *P* < 1e-10), likely due to a relatively constant generation interval in females (σ^2^ = 0.9 years) and a large amount of variation in males across time (σ^2^ = 6.8 years). Males and females reach puberty at approximately the same age (*18*), but the reproductive age in males can extend more than 20 years beyond that in females. Sociocultural factors are likely to have acted in concert with the higher bound on male reproductive age to produce the greater variance observed in male generation interval. The male-female difference follows a similar pattern to that of the average generation time except for the most recent windows, which show a smaller increase in male-female difference than expected during the recent uptick in generation times (compare Fig. 2A and 2B). This smaller difference appears to be driven by a relatively larger increase in recent female generation intervals: the most recent time period is significantly higher than at any point in the last 250,000 years (*P* < 0.005, *z*-test).

To investigate differences in generation times among human populations, we repeated our analysis using four major continental populations within the 1000 Genomes Project. Variants are counted as part of a continental population as long as they are polymorphic among samples from that population. While the continental labels for each population are used across the span of the analysis, note that beyond roughly 2,000 generations ago all non-African populations were likely located in Africa and show little differentiation among themselves; coalescence among all ancestral populations living in Africa does not occur until more than 10,000 generations ago (*15*).

We find subtle changes to the average human generation interval among populations in the last 1,000 generations (Fig. 3; Fig. S7). Average generation times in European and South Asian populations have increased slightly, while generation times in African and East Asian populations have changed little. Similar results in the recent past were observed when using only private alleles (Fig. S8). We estimate a shorter sex-averaged generation interval for Europeans (26.1 years) than East Asians (27.1 years) over the past 40,000 years, supporting a recent estimate derived from divergence to archaic DNA (*11*). Beyond this most recent timeframe, the average generation interval in each of the ancestral non-African populations grows progressively shorter into the past. The dominating pattern across the past 10,000 generations is a significantly shorter sex-averaged generation interval for East Asian, European, and South Asian populations, 20.1 ± 3.9, 20.6 ± 3.8, and 21.0 ± 3.7 years, compared to the African population, 26.9 ± 3.5 years (*P* < 1e-10, *t*-test). The estimated generation times do not converge between populations until we expand our analysis to at least 78,000 generations ago (Fig. 3, inset).

**Figure 3.**
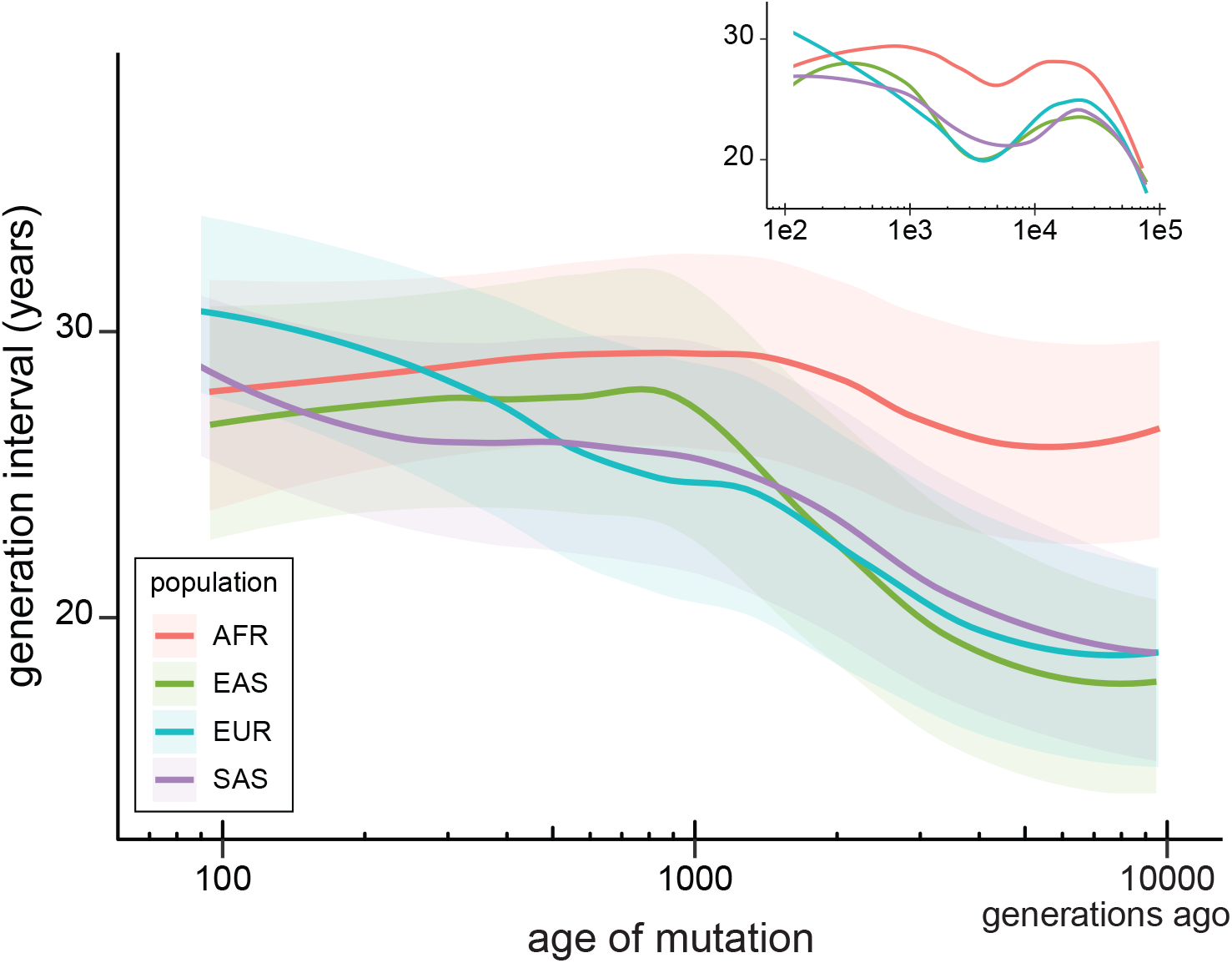
Change in generation interval across different human populations. Generation intervals were estimated in ancestors of four major continental human populations included in the 1000 Genomes Project; sex-averaged generation intervals are shown here as smoothed by loess (see Fig. S7 for full results). Confidence intervals for each population were obtained by bootstrapping, as in Figure 2. The inset shows results from including polymorphisms that date back to 78,000 generations ago, capturing the convergence of estimates in the most recent common ancestor of all populations. AFR: Africa. EAS: East Asia. EUR: Europe. SAS: South Asia.

The large difference in generation times between populations suggests that different timescales are needed to estimate events outside of Africa (20-21 years/generation) versus those in Africa (27 years/generation). These results are consistent with the prediction of a shorter generation time in non-Africans, based on the observation of a slightly elevated per-year mutation rate in these populations (*19*). Note that the difference among populations beyond 2,000 generations ago reflects population structure in humans before their dispersal out of Africa, structure that is not fully captured by the 1000 Genomes AFR sample (*3, 15, 19, 20*). This implies that the simple labels of “African” and “non-African” for these populations conceal differences in generation times that existed on our ancestral continent.

Our study builds upon advances in understanding the characteristics of *de novo* mutations (*14*) and in estimating genome-wide genealogies (*15*) to create a model for generation times that can be applied to ancient populations. While it is clear that the frequency of individual mutation types can evolve rapidly (*16, 17*), even small changefs to the generation interval can reshape the overall mutation spectrum (*21, 22*). Our results are consistent with previous estimates of the average generation time over the past 40-45 thousand years (*10, 11*), but offer unprecedented resolution of sex-specific generation times across 250,000 years of human history. While information on the mutation spectrum far into the past (>10k generations ago) is limited by the coalescent process (and subsequent lack of ancient polymorphisms), fine-scale estimates of generation times from the most recent 100 generations will be possible with larger population samples (cf. *23*). Large enough samples will bring estimates from population genetic data close enough in time to overlap with historical birth records (e.g. *24*). As it stands, our results offer a unique look into the biology of our ancestors and provide a more detailed picture of human demographic history.

## Acknowledgements

We thank J. Raff and A. Bentley for helpful input early in this project, and D. Schrider for comments on the manuscript. This work was supported by the Precision Health Initiative of Indiana University.

## Supplementary Online Materials for

### S1. Modeling the mutation spectrum as a function of parental age

#### S1.1. Data from Icelandic trios

We developed a parental age model for the mutation spectrum based on data from a large study of *de novo* mutations in an Icelandic population (Jónsson et al. 2017). We briefly summarize the findings from this study here as background for the development of our model. The study detected 101,377 single-nucleotide *de novo* mutations from 1,548 trios with known parental ages at conception. In general, they found an increasing number of mutations with both paternal and maternal age, with different rates of increase for different mutation classes. The parent-of-origin was determined for a subset of these mutations (*n* = 41,899), allowing inferences for the mutation spectrum to be made separately for mothers and fathers. Figure S1 summarizes these findings for each of the six different classes of single-nucleotide mutations (A→C, A→G, A→T, C→A, C→G, C→T; each class includes counts from their complements).

**Figure S1.**
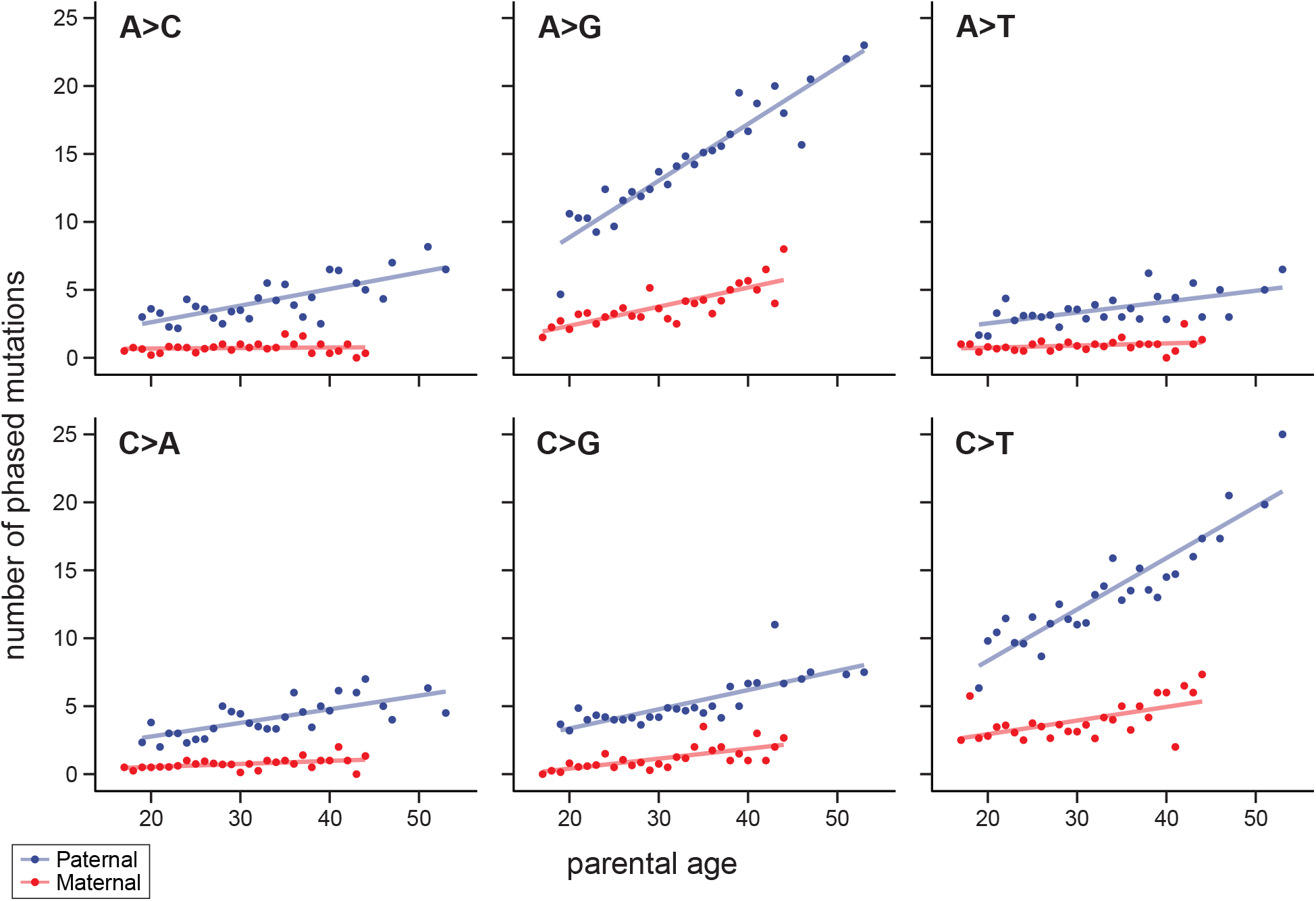
Frequency of mutation classes with parental age. A summary of the number of *de novo* mutations as a function of age. Phased mutations can be assigned to either the paternal or maternal lineage, so are shown separately for the six different types of single nucleotide changes (and their complement). Data from Jónsson et al. (2017).

#### S1.2. Description of the Dirichlet-multinomial regression

The mutation spectrum is a form of compositional data: comparisons between spectra focus on differences in the relative abundance of each mutation class. Because of the small number of mutations produced by any one set of parents, observations from a single trio are insufficient to reliably determine the spectrum. A model for the mutation spectrum must therefore incorporate the probabilistic nature of mutation counts from a given trio while inferring the relationship between the underlying spectrum and given covariates. We apply a Dirichlet-multinomial regression to mutation count data to capture the relationship between the underlying mutation spectrum and parental ages, which are treated as covariates in the analysis.

Let ***y***_*i*_ = (*y*_*i*,A→C_, *y*_*i*,A→G_, *y*_*i*,A→T_, *y*_*i*,C→A_, *y*_*i*,C→G_, *y*_*i*,C→T_ be the vector of mutation counts for each of the six respective mutation classes from trio *i*. The distribution for *m* mutation counts from a trio, **y**_*i*_, is modeled as a multinomial, conditional on the probability vector ***p_i_***,

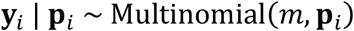

where ***p***_*i*_ is defined on the 6-dimensional simplex, *S* = {(*p*_A→C_, *p*_A→G_, *p*_A→T_, *p*_C→A_, *p*_C→G_, *p*_C→T_): *P_j_* ≥ 0, ∑_*j*_ *p_j_* = 1}.

We then impose a conjugate Dirichlet prior on **p**, such that **p** ~ Dirichlet(**α**), and **α** = (α_A→C_, α_A→G_, α_A→T_, α_C→A_, α_C→G_, α_C→T_), α_*j*_ > 0. The probability mass function for the count vector **y** over *m* = **∑**_*i*_ ***y**_i_* trials under this Dirichlet-multinomial model can be represented as

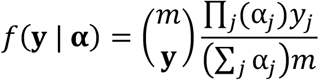

(see Kim et al. 2018).

The parental ages for each trio are incorporated as covariates for the Dirichlet-multinomial regression, **x** = (**x**_paternal_, **x**_maternal_, an *n* × 2 matrix of parental ages. They are related to the Dirichlet parameter **α** by the inverse link function,

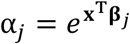

where **β**_*j*_ = (*β*_*j*, paternal_, *β*_*j*, maternal_) is the vector of regression coefficients for each mutation class.

#### S1.3. Subset of mutations and trios for model fitting

For our main analysis we used a subset of mutations from the Icelandic dataset to model the mutation spectrum with parental age: we used only the set of phased mutations for which the parent-of-origin was determined by either read-tracing or transmission to a third generation. Further restrictions on the mutations used for modeling were made to mirror the filters placed on dated variants from the 1000 Genomes Project dataset. These include removing mutations at CpG sites and C→T transitions with a trinucleotide context associated with a putative mutation pulse (see section *S2.2*). We also restricted trios to those that had a minimum of at least 10 mutations. This was done to avoid matrix degeneracy when fitting the maximum likelihood mutation spectrum model (see below). After all filters, we fit the model on 27,902 mutations from 1,247 trios.

#### S1.4. Fitting the model to mutation data

We used the R package MGLM (Kim et al. 2018) to fit the Dirichlet-multinomial regression model to the filtered mutation dataset. MGLM implements several methods for multivariate generalized linear models, including the Dirichlet-multinomial. We used it to fit the regression coefficients for our covariates (parental age) that maximize the log-likelihood of our model. The result is a predictive model that gives the expected mutation spectrum for a set of parental ages. Figure S2 demonstrates a set of simple predictions from the fit model, showing the expected changes to the mutation spectrum when paternal and maternal age are individually adjusted.

**Figure S2.**
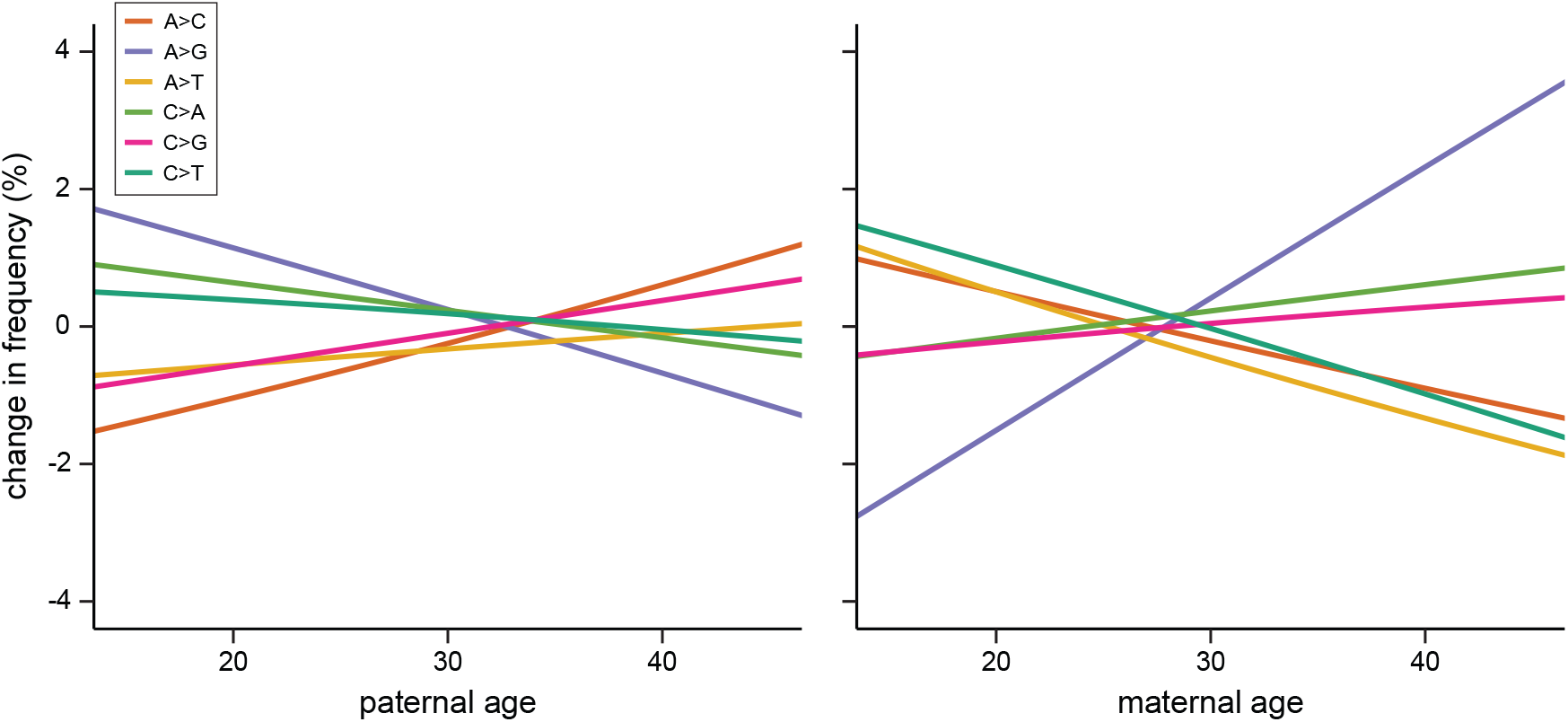
Predicted change in mutation frequency with paternal and maternal age. Data from Icelandic trios (Figure S1) were used to parameterize the Dirichlet-multinomial model. Figures are centered on the average paternal and maternal ages among the trios, so that predicted changes in each type of mutation can be visualized as a change relative to this age.

### S2. Variants from the 1000 Genomes Project dated by GEVA

#### S2.1. GEVA and the Atlas of Variant Age

Human variants dated by the Genealogical Estimation of Variant Age (GEVA) approach are publicly available in the Atlas of Variant Age, an online database (https://human.genome.dating/). We used dated variants in this database collected from the 1000 Genomes Project (Phase 3; GRCh37). This set includes autosomal variants sampled from 2,504 individuals in 26 worldwide subpopulations within 5 continental populations. Ancestral and derived states were determined in the Atlas of Variant Age through multispecies alignments from the Ensembl database (see Albers and McVean 2020). Throughout our analysis, we use the median estimated allele age from the database as a point estimate of each variant’s age.

#### S2.2. Filtering dated variants

We took several additional filtering steps to ensure variants were appropriate for estimating generation time. We considered only biallelic single-nucleotide sites that were not singletons— variants that exist on only a single chromosome across samples. We also discarded variants with a derived allele frequency higher than 98% to reduce the possibility of ancestral state misidentification.

CpG sites are more likely to have arisen more than once, and therefore to have been multiply mapped on genealogies; their frequency is much less consistent across time periods as a result (Fig. S3). As in the model for mutation spectrum with parental age, all variants at CpG sites were discarded from consideration.

Several C→T transitions have been inferred to be part of a recent mutation pulse, particularly in European populations (Harris and Pritchard 2017). To reduce the potential effect of this mutation pulse on estimates of generation time, we discarded all triplet C→T transitions that have been found to be associated with this pulse. These include ACC→ATC, CCC→CTC, TCC→TTC, TCT→TTT, and their respective reverse complements.

#### S2.3. Binning data into time periods

After all filtering, there were 25.3 million variants from the Atlas of Variant Age for which there were estimates of allele age. Of these, 20.9 million were estimated to have arisen in the last 10,000 generations. Because there are very few young variants and a long tail for the number of older variants (Fig. S4), we estimated spectra in bins that were supported by equal numbers of variants rather than in equally spaced time periods. We divided the 20.9 million variants equally among 100 bins based on their estimated age. Bins were filled starting with the youngest variants, leaving a small number of the remainder of oldest variants unplaced.

The estimated spectrum for each bin was calculated as the count of variants in each of the six mutation classes divided by the total number of variants in the bin. The age of each bin was calculated as the mean of estimated ages from all variants in the bin. Figure S3 shows the spectra, as a frequency of each mutation class, across 100 bins from the past 10,000 generations. The same procedure was used to estimate historic spectra for each of the continental population groups, for which there were 11.0 (AFR), 4.3 (EAS), 4.4 (EUR), and 5.4 (SAS) million variants included after filtering (see section *S4.1*).

**Figure S3.**
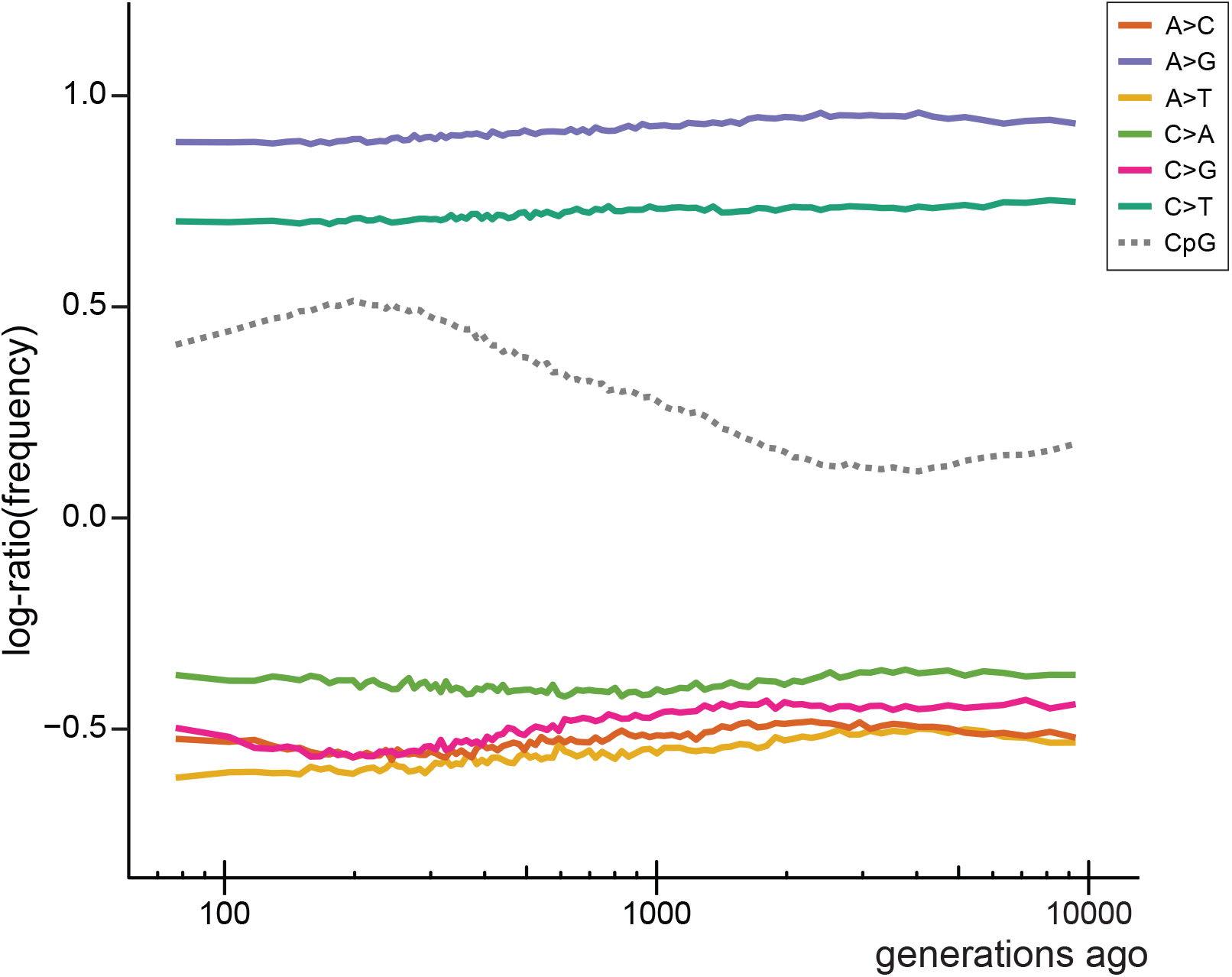
Mutation frequency by age of origin. For each of 100 time periods, the frequency of each type of mutation having been inferred to arise in that bin is plotted. In addition to the six types of mutations used in the Dirichlet-multinomial model, we also show the behavior of CpG→CpT mutations for comparison (these were not used in the model).

**Figure S4.**
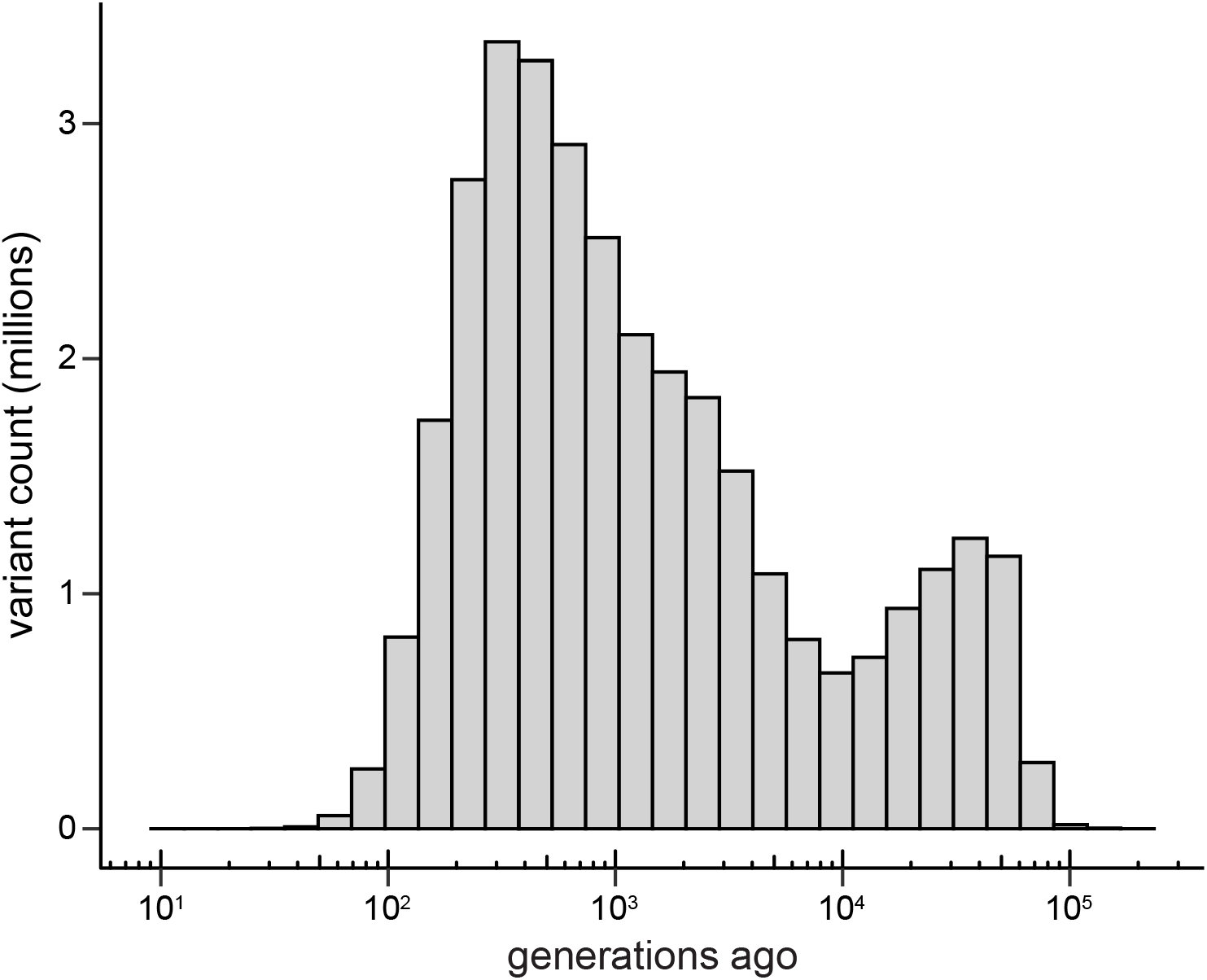
Density of variants by age of origin. Variants dated by GEVA (Albers and McVean 2020) are plotted according to the time at which they are estimated to have arisen via mutation. The plot includes all data from the 1000 Genomes Project, regardless of which population(s) they are found in.

### S3. Estimating generation times

#### S3.1. Fitting variant data to the Dirichlet-multinomial regression model

We jointly estimate separate male and female generation times from the historic mutation spectra calculated from the counts of variants in each time period. To do this, the parental ages in the Dirichlet-multinomial model were treated as parameters in a search for a predicted mutation spectrum that best fit the observed historic spectrum. We minimized the distance between each predicted spectrum and each observed historic spectrum.

Because a mutation spectrum is a composition underlain by count data, comparisons between spectra using simple Euclidean distance can be misleading. Like all compositional data, mutation spectra are mathematically constrained by the possible values for the frequency of each count class, distorting the simple Euclidean distance between compositions. To deal with this, we perform a centered log-ratio transformation (clr) on each spectrum before calculating the distance between them (Aitchison 1986). The transformation can be obtained as

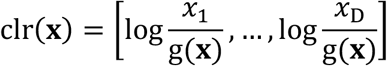

for a composition vector **x** with D elements, where g(**x**) is the geometric mean of the composition. The Aitchison distance between two given spectra, **x**_1_ and **x**_2_, can then be calculated as *d* = ∥clr(**x**_1_) – clr(**x**_2_)∥.

We found that the mutation spectrum from the Icelandic trios (Jónsson et al. 2017) differs from the variant spectrum inferred from the 1000 Genomes Project data across all time periods. This difference is relatively constant across the time periods we considered. Therefore, to obtain absolute generation times for historic periods, we center the observed spectra on the most recent bin, subtracting the difference between it and the average mutation spectrum estimated in Jónsson et al. (2017) from each historic spectrum. This has the effect of assuming that parental ages in the Icelandic dataset reflect generation times in the most recent historic bin. We find this assumption to be robust for both the relative difference in generation time between the sexes as well as the overall pattern of historic generation times (see section *S3.4*).

The generation time was then estimated from each historic mutation spectra by distance minimization as

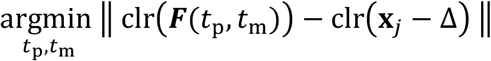

where ***F*** gives the predicted spectrum from the Dirichlet-multinomial model for a set of paternal and maternal ages, *t*_p_ and *t*_m_, **x**_*j*_ is the historic mutation spectrum from a given time period, and Δ is the centering difference described above. The parental ages that minimized this distance were found by applying the L-BFGS-B optimization algorithm as implemented in the R *stats* package (Nocedal and Wright 1999). We used the default convergence tolerance, maximum number of iterations, and set bounds for both parental ages to be: [0, 100]. None of the searches returned a minimum distance at these bounds. The maternal and paternal ages that minimized the distance from each time period were taken to be the respective estimates of the generation time. These ages, as well as the sex-averaged generation time, for all time periods are provided in Supplementary Data File 1.

#### S3.2. Calculating confidence intervals by double-bootstrap

There are two major sources of uncertainty in our estimates of the generation time: (1) the mutation data that specifies the Dirichlet-multinomial regression model, and (2) the dated variants that are used to calculate the variant spectrum in each time period. This led us to construct confidence intervals around the generation time estimates with a double-bootstrap resampling strategy.

The 1,247 trios from the Icelandic dataset were resampled with replacement and fit to the Dirichlet-multinomial regression model. We discarded cases where the likelihood search for the regression model failed to converge, but restricting the dataset to include only trios that had at least 10 mutations greatly reduced instances of failure to converge due to matrix singularity. The variants in each time period of the analysis were also resampled with replacement and the spectrum was recalculated for each bin. Finally, generation times were estimated by fitting the bootstrapped spectrum to the bootstrapped model by distance minimization as described above. The resampling steps were each repeated 100 times, resulting in a total 100 × 100 = 10,000 bootstrap estimates of generation time for each time period included in the analysis.

#### S3.3. Calculating averages and absolute generation times

The sex-averaged generation time was calculated as the mean of the maternal and paternal ages estimated for each time period. In figures plotting this sex-averaged estimate, we performed local polynomial regression (loess) to produce a smoothed curve across the past. We used the default smoothing parameter, α = 0.75, in the R *stats* implementation of loess to smooth both sex-averaged estimates and their confidence intervals.

We calculated the absolute time scale (Fig 2A in main text) on which generation times change by integrating the estimated sex-averaged generation time across the age of mutations. We employed a Riemann sum, calculating the cumulative sum of estimated generation times in single generation steps from the smoothed sex-averaged curve. We added a small constant to this integration to account for the time between the present and the first estimate by assuming there has been no change to generation times in this short period.

A related strategy was used to calculate the average generation times across the period of our analysis. Because ranges for each time period were based on an equal number of variants, older bins span a greater amount of time. We weighted the estimate from each time period by the span of the bin when calculating the average generation times reported in the main text.

#### S3.4. Limited effect of recombination on generation time estimates

Recombination could distort our generation time estimates if linked selection or biased gene conversion affects the inferred date of origin of variants in a way that nonuniformly changes historic spectra. Linked selection will change the shape of genealogies (Barton 1998), especially in regions of low recombination. GC-biased gene conversion will change the population frequency of specific variants, but has a greater effect in regions of high recombination (Lachance and Tishkoff 2014; Glémin et al. 2015).

We did not expect either process to affect the dating of variants within genealogies, but carried out additional analyses to ensure the robustness of our results. We assessed whether recombination might have affected generation time estimates by repeating our analysis with dated variants from regions with different recombination rates in the human genome. We split variants into quintiles based on the interpolated human map of recombination (https://github.com/joepickrell/1000-genomes-genetic-maps). While our estimates of generation time appear to show a slight increase with increasing recombination (Fig. S5A), the pattern across history remains the same and estimates from all quintiles fall within the bootstrapped confidence intervals. This slight increase seen in estimates from regions of higher recombination did not correspond with any bias toward G or C alleles in the inferred mutation spectra from these regions (Fig. S5B). It is therefore not clear how recombination leads to the small observed changes in generation time estimates.

**Figure S5.**
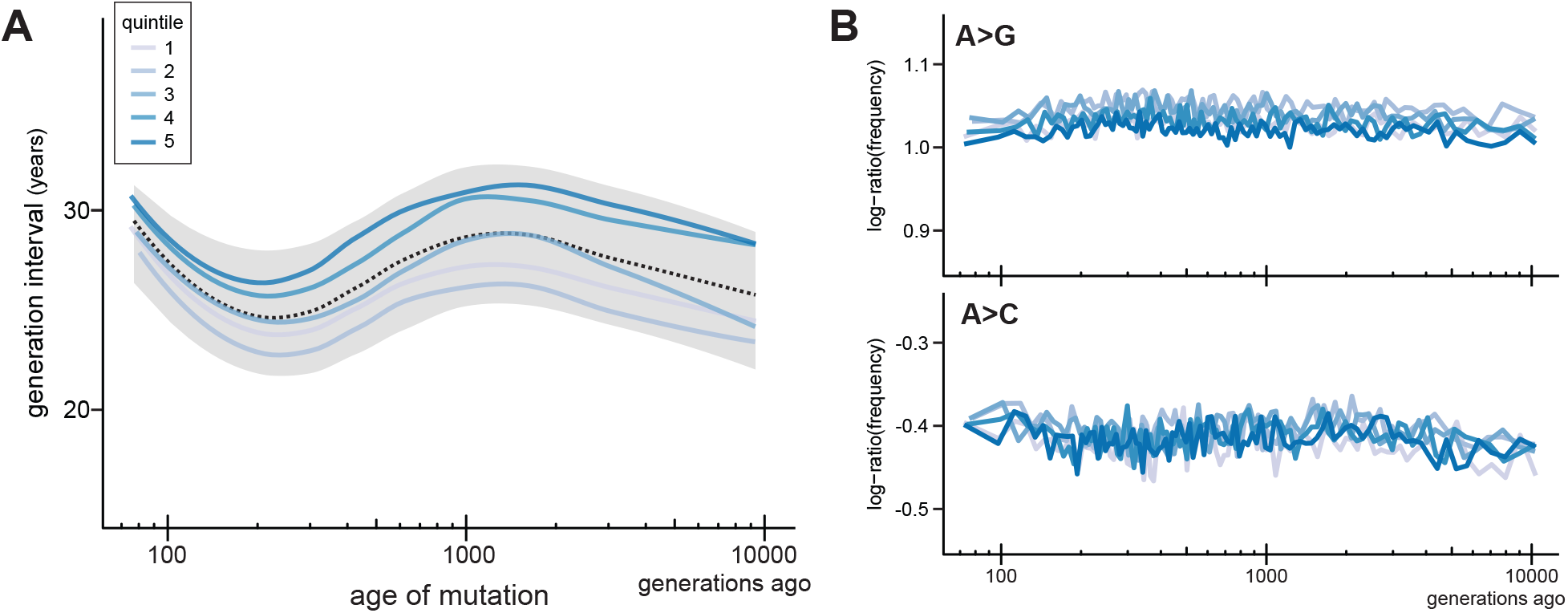
Generation intervals estimated from variants with different recombination rates. A) Inferences of sex-averaged generation times for data taken from variants binned according to the local recombination rate. Variants were binned into five quintiles of recombination rate, ranging from the lowest recombination rate regions (quintile 1) to the highest recombination rate regions (quintile 5). B) Change in the frequency of mutations to G and C over time, binned by local recombination rate.

#### S3.5. Additional effects of relaxing filters and assumptions

We examined several ways in which data or modeling choices might have affected our results. Rather than using only the set of high-quality phased mutations, we fit the Dirichlet-multinomial regression model to a much larger dataset that included unphased mutations from the Icelandic trios (*n* = 72,573 *de novo* mutations). The results from this analysis are shown in Figure S6A. The male-female difference is slightly accentuated, but the overall pattern for generation times remains the same.

As mentioned in section *S3.1*, the main results were anchored by absolute generation time estimates from the most recent time period. We relaxed this assumption by anchoring to the mean spectrum across all dated variants. This effectively asserts that the Icelandic dataset has a generation time equivalent to the mean generation time across thousands of generations. While estimates of absolute parental age were slightly lower under this assumption, the patterns across human history were unaffected (Fig. S6B).

**Figure S6.**
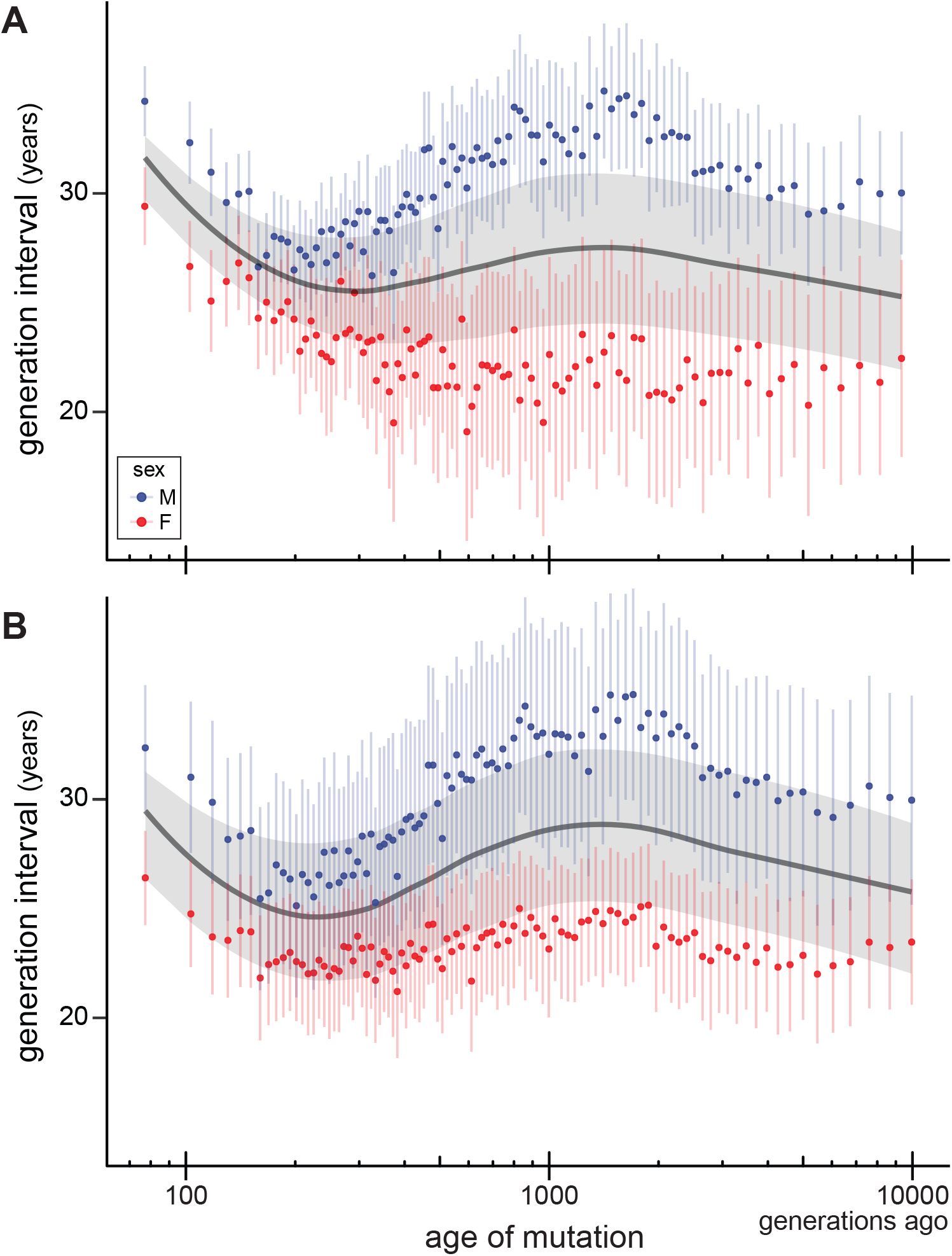
Generation intervals estimated from model that includes unphased mutations. A) All *de novo* mutations from the Icelandic trio dataset (not just phased mutations, as in Fig. 2 in main text) were used to re-parameterize the Dirichlet-multinomial model, and then to reestimate generation times. B) Generation times estimated by anchoring the Icelandic mutation frequency spectrum to the average frequency spectrum across all historic time periods.

### S4. Estimating population-specific generation times

#### S4.1. Separating the 1000 Genomes Project data into continental populations

We separated variants as belonging to one of four continental populations (AFR: Africa, EAS: East Asia, EUR: Europe, and SAS: Southeast Asia) based on their geographic sampling in the 1000 Genomes Project. Variants were placed into continental populations using an inclusive criterion: as long as more than one copy exists among samples from a population, it is included in that population. We analyzed each set of variants separately to arrive at population-specific estimates of generation times (Fig. S7). That is, we repeated each step of the previously described analysis with only the subset of variants included in each population.

**Figure S7.**
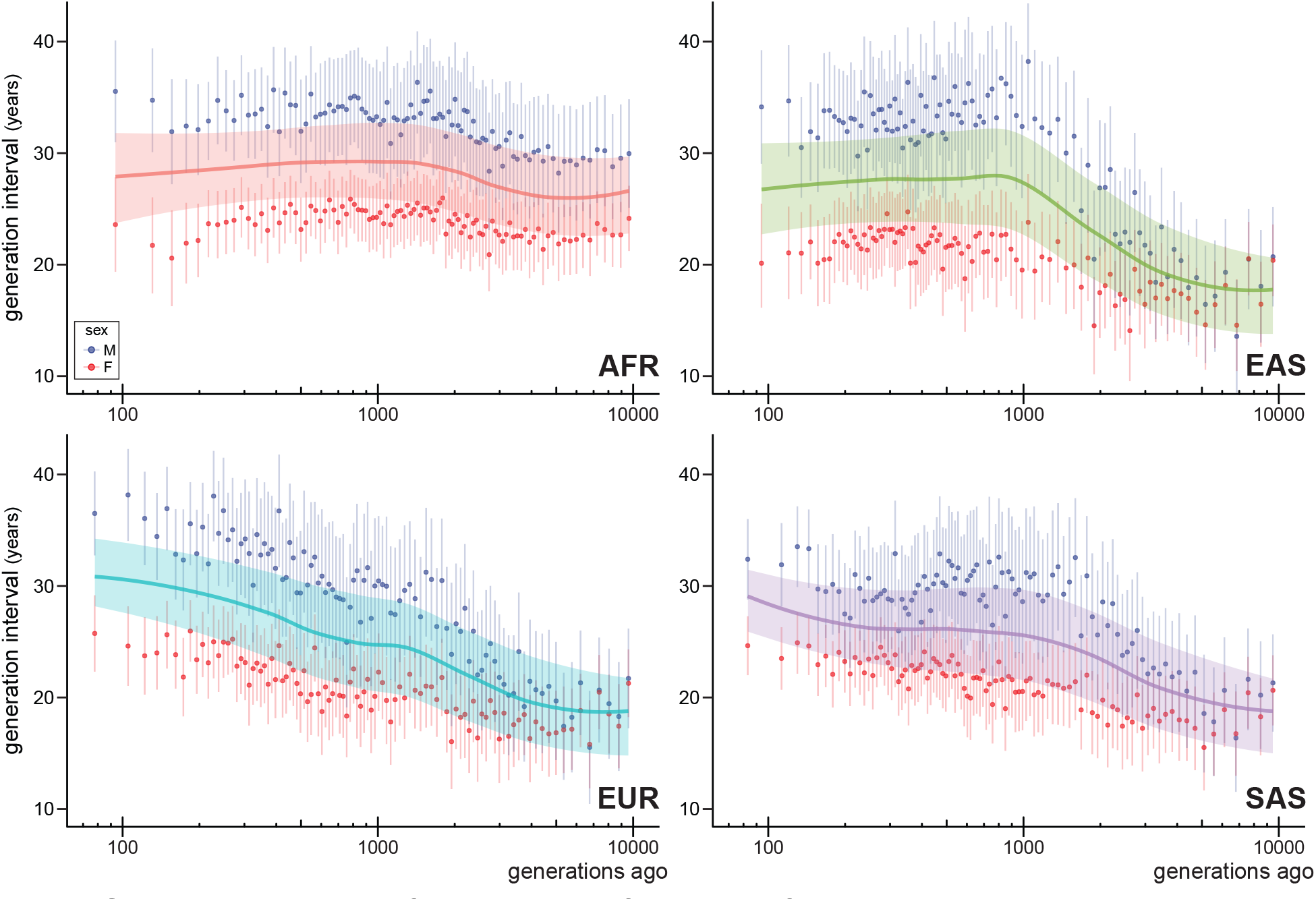
Population-specific estimates of male and female generation interval. Generation intervals were estimated for four major continental populations. These results are the same as those shown in Figure 3 in the main text, but with separate maternal and paternal generation times plotted.

#### S4.2. Estimates from private alleles

In contrast to the broadly inclusive criteria, we also separated variants into each continental population by including only the private alleles exclusive to each population. The proportion of variants that are private to each continental population drops rapidly going back in time, and they make up a very small proportion of variants by 2,000 generations ago (Fig. S8B). Nevertheless, we estimated generation times after creating a subset of variants for each population using only the private alleles. Figure S8A shows the results of this analysis for the first 1,000 generations, before private variants for most populations disappear. These results are very similar to those found using the more inclusive criteria for variants (Fig. 3 in the main text).

**Figure S8.**
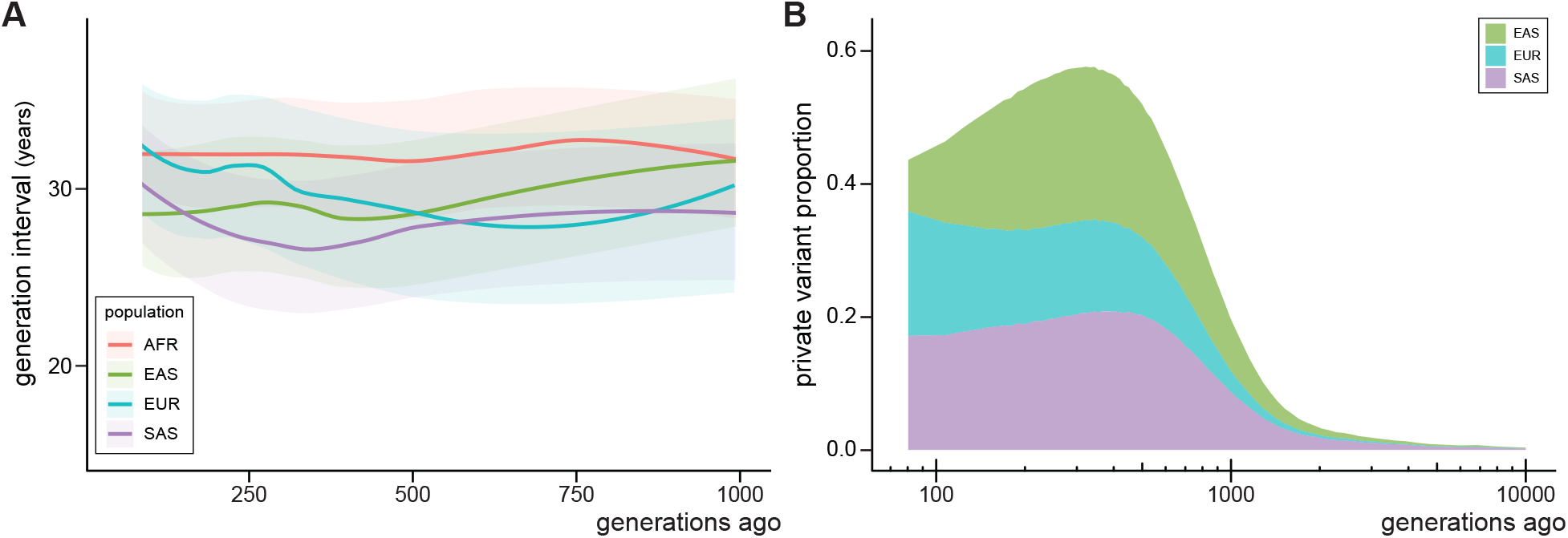
Population-specific estimates from private variants. A) Estimates of the generation interval for each of the four major continental populations using only variants private to each population. These results can be compared to Figure 3 in the main text, but note that here we only plot estimates up to 1000 generations ago. B) The proportion of all variation that is private to one continental population, as a function of time in the past. Almost all variation private to one of the non-African samples has arisen in the most recent 1000 generations.

## References

1. H. Li, R. Durbin, Inference of human population history from individual whole-genome sequences. Nature 475, 493–496 (2011).

2. L. Speidel, M. Forest, S. Shi, S. R. Myers, A method for genome-wide genealogy estimation for thousands of samples. Nature Genetics 51, 1321–1329 (2019).

3. A.-S. Malaspinas et al., A genomic history of Aboriginal Australia. Nature 538, 207–214 (2016).

4. E. Huerta-Sánchez et al., Altitude adaptation in Tibetans caused by introgression of Denisovan-like DNA. Nature 512, 194–197 (2014).

5. J. N. Fenner, Cross-cultural estimation of the human generation interval for use in genetics-based population divergence studies. American Journal of Physical Anthropology 128, 415–423 (2005).

6. S. Matsumura, P. Forster, Generation time and effective population size in Polar Eskimos. Proceedings of the Royal Society B: Biological Sciences 275, 1501–1508 (2008).

7. J. P. Bocquet-Appel, When the world’s population took off: the springboard of the Neolithic Demographic Transition. Science 333, 560–561 (2011).

8. P. Jordan, in The Oxford Handbook of the Archaeology and Anthropology of Hunter-Gatherers, V. Cummings, P. Jordan, M. Zvelebil, Eds. (Oxford University Press, Oxford, 2014).

9. K. E. Langergraber et al., Generation times in wild chimpanzees and gorillas suggest earlier divergence times in great ape and human evolution. Proceedings of the National Academy of Sciences 109, 15716–15721 (2012).

10. P. Moorjani et al., A genetic method for dating ancient genomes provides a direct estimate of human generation interval in the last 45,000 years. Proceedings of the National Academy of Sciences 113, 5652–5657 (2016).

11. M. Coll Macià, L. Skov, B. M. Peter, M. H. Schierup, Different historical generation intervals in human populations inferred from Neanderthal fragment lengths and mutation signatures. Nature Communications 12, 5317 (2021).

12. A. Kong et al., Rate of *de novo* mutations and the importance of father’s age to disease risk. Nature 488, 471–475 (2012).

13. R. Rahbari et al., Timing, rates and spectra of human germline mutation. Nature Genetics 48, 126–133 (2016).

14. H. Jónsson et al., Parental influence on human germline *de novo* mutations in 1,548 trios from Iceland. Nature 549, 519–522 (2017).

15. P. K. Albers, G. McVean, Dating genomic variants and shared ancestry in populationscale sequencing data. PLoS Biology 18, e3000586 (2020).

16. K. Harris, Evidence for recent, population-specific evolution of the human mutation rate. Proceedings of the National Academy of Sciences 112, 3439–3444 (2015).

17. K. Harris, J. K. Pritchard, Rapid evolution of the human mutation spectrum. eLife 6, e24284 (2017).

18. A.-S. Parent et al., The timing of normal puberty and the age limits of sexual precocity: variations around the world, secular trends, and changes after migration. Endocrine Reviews 24, 668–693 (2003).

19. S. Mallick et al., The Simons Genome Diversity Project: 300 genomes from 142 diverse populations. Nature 538, 201–206 (2016).

20. E. M. Scerri et al., Did our species evolve in subdivided populations across Africa, and why does it matter? Trends in Ecology & Evolution 33, 582–594 (2018).

21. J. Carlson, W. S. DeWitt, K. Harris, Inferring evolutionary dynamics of mutation rates through the lens of mutation spectrum variation. Current Opinion in Genetics & Development 62, 50–57 (2020).

22. R. J. Wang et al., De novo mutations in domestic cat are consistent with an effect of reproductive longevity on both the rate and spectrum of mutations. bioRxiv, 2021.2004.2006.438608 (2021).

23. A. Keinan, A. G. Clark, Recent explosive human population growth has resulted in an excess of rare genetic variants. Science 336, 740–743 (2012).

24. A. Helgason, B. Hrafnkelsson, J. R. Gulcher, R. Ward, K. Stefánsson, A populationwide coalescent analysis of Icelandic matrilineal and patrilineal genealogies: evidence for a faster evolutionary rate of mtDNA lineages than Y chromosomes. The American Journal of Human Genetics 72, 1370–1388 (2003).

## Supplementary References

Albers, P. K., and G. McVean. 2020. Dating genomic variants and shared ancestry in population-scale sequencing data. PLoS Biology 18:e3000586.

Aitchison, J. 1986. The statistical analysis of compositional data. Chapman and Hall, London-New York.

Barton, N. H. 1998. The effect of hitch-hiking on neutral genealogies. Genetical Research 72:123–133.

Glémin, S., P. F. Arndt, P. W. Messer, D. Petrov, N. Galtier, and L. Duret. 2015. Quantification of GC-biased gene conversion in the human genome. Genome Research 25:1215–1228.

Harris, K., and J. K. Pritchard. 2017. Rapid evolution of the human mutation spectrum. eLife 6:e24284.

Jónsson, H., P. Sulem, B. Kehr et al. 2017. Parental influence on human germline *de novo* mutations in 1,548 trios from Iceland. Nature 549:519–522.

Kim, J., Y. Zhang, J. Day, and H. Zhou. 2018. MGLM: An R package for multivariate categorical data analysis. The R Journal 10:73–90.

Lachance, J., and S. A. Tishkoff. 2014. Biased gene conversion skews allele frequencies in human populations, increasing the disease burden of recessive alleles. American Journal of Human Genetics 95:408–420.

Nocedal, J., and S. J. Wright. 1999. Numerical optimization. Springer, New York.

